# Mechanism of Membrane Recovery in Intra-Cytoplasmic Sperm Injection

**DOI:** 10.1101/240200

**Authors:** Ricky Li, Hao Chang, Brian Luo

## Abstract

ICSI (Intra-cytoplasmic sperm injection) is a broadly utilized technique for artificial fertilization. This approach has been successfully performed in human oocytes as well as others such as mouse and bovine. The piercing through the zona layer and the membrane needs to be achieved with a minimal biological damage to facilitate a rapid healing. Since the injection methodology serves as a crucial factor to success rate of ICSI, a significant amount of research efforts has been devoted to the development of injections. In this paper, we conduct comparative study among the major milestones for injection techniques in ICSI. Technical details are provided for each milestone and each technique is evaluated from engineering perspective. Later, we present a mechanism for healing process of membrane after drilling, which could potentially provide guidance for improvement of injection method. More importantly, we perform coarse-grained molecular dynamics simulation to reveal the mechanism of membrane recovery in intra-cytoplasmic sperm injection.

## 1. Introduction

As a broadly utilized technique for artificial fertilization, ICSI (Intra-cytoplasmic sperm injection) has been successfully performed in human oocytes as well as others such as mouse and bovine [1-6]. Among the applications of ICSI, fertilization in mouse is of particular interest to biological and biomedical research. The significance of biological cell injection technology has attracted large amounts of research in an effort to automate laborious cell injection tasks. For the previous decade, many research efforts are dedicated to cell injection automation from a diverse array of aspects: cell holding devices, cell injection method, vision-based automation, cell injection process control, etc [7-8]. Conventional ICSI employs a spiked micropipette for facilitating penetration of the zona pellucida and piercing of the oolemma to enable penetration into the ooplasm and injection of whole sperm into the ovum. However, the irreversible damage to the ovum can be caused by the elasticity of the mouse oolemma, which makes it difficult to penetrate without rupture [9-10,63]. A minimal biological damage has to be achieved during the piercing through the zona layer and the membrane to facilitate a rapid healing [9]. A significant amount of research effort has been devoted to the development of ICSI from perspective of injection technique.

In conventional cellular injection for ICSI as shown in Fig. 1, an individual oocyte is immobilized by a holding pipette with a slight suction. The injection pipette containing the sperm head to be injected into the oocyte is forced into the cell. The survival rates and fertilization rates of oocytes for conventional ICSI vary from 80 to 90%, and 45 to 70%, respectively [11-16]. Size and sharpness of the needle used for injection are reported to be important for success rate and among the crucial factors is injection method of the needle into ooplasm [14].

**Fig. 1:**
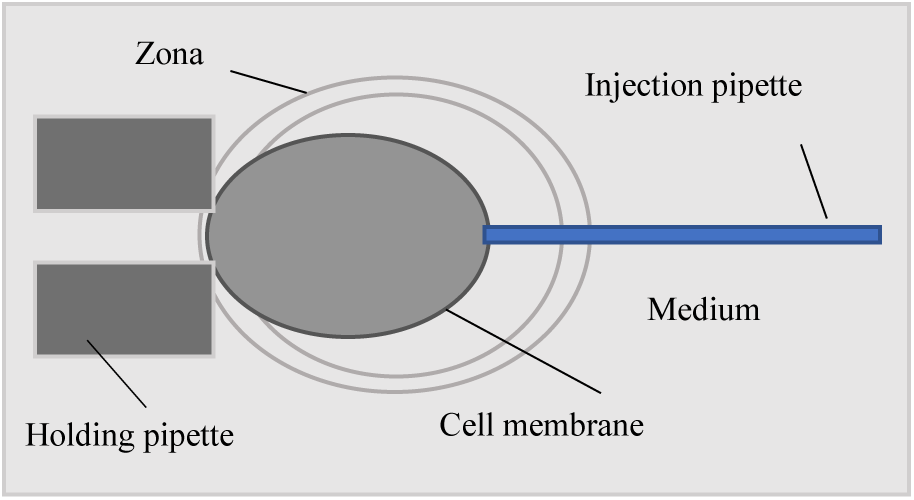
Intra-cytoplasmic sperm injection setup.

Due to the low success rate and high labor intensity, piezo-ICSI was firstly employed to mice in 1995 [25-30,64-66], which achieved high survival and fertilization rates in mice compared with those obtained by conventional ICSI. Despite the noticeably higher efficiency than conventional ICSI, in current practice, piezo-assisted ICSI requires the presence of a small amount of mercury to stabilize and suppress undesired lateral vibrations of the pipette tip under mechanical impact [17-20]. As of today, no other fluid material can replace mercury which can provide sufficiently high density necessary to mitigate these effects, preventing injection-induced damage to the ovum. Unfortunately, massive deployment of piezo-assisted ICSI in labs and institutions is prevented by the toxicity of mercury. Over the years, researchers have been taking great efforts to find alternative injection method to eliminate dependence on toxic substance in ICSI.

Later, a novel, mercury-free, rotationally oscillating drill (Ros-Drill) device was proposed as an alternative to piezo-assisted ICSI. Since it doesn’t employ piezoelectric force actuator to drive a micropipette for performing ICSI in the mouse, it does not require mercury to suppress unwanted lateral motion of piezo. Preliminary results from Ros-Drill in mouse embryos was reported to be comparable to those of piezo-assisted ICSI using mercury. Moreover, minimal demand on human expertise was claimed to be another benefit of this technology. These features result from the computer automated nature of the Ros-Drill technology as explained in the recent technology paper [21-23]. However, there is very few experimental or clinic data to support the statements and successful duplication by this method is necessary.

Most recent improvements in ICSI revolve around piezo-ICSI. Two main directions for improvement are micropipettes design to improve fertilization rate and injector design to avoid usage of mercury. For example, a piezoelectric driven non-toxic injector for automated cell manipulation was proposed [24], which claims to suppress detrimental lateral tip oscillation by centralizing the piezo oscillation power on the injector pipette However, so far only pipette movement is captured under high speed camera to show attenuation of lateral movement. Experimental and clinic data is not available now. Capability of piezo-ICSI may be compromised due to limiter applied in lateral dimension.

In this paper, at first, comparative study is presented for the major milestones for injection techniques in ICSI from extensive aspects. In essence, understanding of mechanism of membrane healing is crucial in ICSI from two-folds: a. new injection method to facilitate healing of membrane, b. facilitate healing after injection. At last, we present a mechanism for membrane healing process after injection.

## 2. Major milestones

In this section, technical details are provided for each milestone and each technique is evaluated from engineering perspective.

### 2.1 Conventional ICSI

The first method for ICSI is performed manually and is known as conventional ICSI [25-35]. With this technique, the tip of the injection pipette is approaching gently towards the cell membrane about halfway or even much further into the oocyte. Then it is pushed against membrane rapidly until it penetrates through the membrane. The spike-shaped pipette is most commonly used for conventional ICSI [26]. The high compliance of the mouse membrane makes the penetration very difficult without rupturing the ovum [14]. High failure rate for conventional ICSI can be attributed to the damage caused by the pressure of the injecting pipette on the cell membrane during the piercing process. Damage needs to heal properly and rapidly; otherwise abnormal growth occurs in the future stages of development [14,32,44,67]. Since manual operation can’t perform piercing process quickly and accurately enough, deformation induced pressure inside membrane inevitably accumulates during piercing process. In light of limitation in manual operation, injection parameters such as injection depth (deformation level) and contact point become more critical and stringent, which requires extensive study in membrane modeling and intensive training in biologists and practitioners.

### 2.2 Piezo-assisted ICSI

The most popular procedure at the present is the piezo-assisted ICSI which has proved to increase the success rate beyond the conventional ICSI results [36-45]. In this technique, to generate minimal damage during the injection process, a piezo-activated axial force is applied to the injecting pipette to pierce smoothly the zona pellucida and then the cell membrane. Different from conventional ICSI, for piezo-assisted ICSI penetration of the zona pellucida and oolemma can be facilitated with a flat-tipped (not spiked) micropipette [38]. In piezo-actuated micromanipulation, the piezo-electric effect is employed to transmit a small crystal lattice distortion to the tip of a pipette, moving it forward against membrane in a prescribed manner. Piezo-actuated micromanipulation has multiple applications in the study and engineering of gametes and embryos. For example, it enabled the first intracytoplasmic sperm injection (ICSI) to produce mice, the first nuclear transfer cloning of mice and pigs [41-42].

The typical controller parameters of piezo-assisted ICSI are set to 10 V amplitude, 60s duration, and 2 Hz frequency for mouse oocytes [36,46-48]. Introduction of piezo-actuation into ICSI for precise and consistent control is, in fact, a significant progress towards an effective automated deployment of micro-injection operation. However, a small amount of mercury is usually placed near the tip of piezo-actuated pipette to suppress its transverse oscillations [17,18,42]. This high intensity substance can effective suppress vibration as the damping effect in lateral dimension.

The major supplier for Piezo-actuated micromanipulation are Eppendorf PiezoXpert and Burleigh© Piezodrill. Usually, their devices enable tuning of parameters, such as speed and number of pulse and ramp-up/down slope for pulse [46,47]. These parameters need to be optimized after collections of experiments are conducted to achieve good results.

### 2.3 Ros-Drill

The main operational difference between piezo-drill and Ros-Drill is in the motion generation methods. Instead of the axial impact type action in the piezo-drill, the Ros-Drill rotationally oscillates the micropipette at a selected frequency and amplitude [21-22]. The intention for Ros-Drill design is that the ideal straight micropipette would rotationally oscillate about its axis without a lateral motion. However, the eccentricity in pipette due to manufacturing tolerance would generate some whirling motion during oscillation [21-22]. Therefore, the eccentricity level determines the unwanted lateral movement.

In alpha version of Ros-Drill, to avoid excessive lateral displacements, pipette is oscillated with very small angular amplitude (1° peak-to-peak) and at frequencies that above natural frequencies of mode 1 and mode 2 (90–100 Hz) and much lower than the 3^rd^ mode [21]. 30° bent pipettes are selected for during injection. This Ros-Drill microinjector prototype employs a PLC (Programmable Logic Controller) as the digital controller, which has a maximum sampling frequency of 1 kHz [22]. This constraint limits the controllable trajectory to be at the maximum frequency of 500 Hz due to the Nyquist sampling rule. A DC servo motor is employed to actuate the pipette to track a sinusoidal position reference. The servo motor comes with an optical encoder with 512 lines and quadrature signature capability which could only provide a sensor resolution of 0.176° [22].

Most recently reported ICSI tests use injection pipette oscillations up to 0.2° amplitude and maximum frequency of 500 Hz [22]. Small angular oscillation is to minimize excessive lateral motion. High frequency is adopted to avoid cell membrane to follow the stimulus, facilitating a clean piercing process. One assumption is made in this new technique that the motion generated from piezo-actuator is faithfully transmitted to tip of pipette due to high rotational stiffness of the pipette holder and the pipette. Experiments were done to show that Ros-Drill operation generates smaller lateral oscillations at the tip, compared with the piezo-assisted ICSI cases [21-22]. Although biological results based on this prototype are very promising, the trajectory tracking performance of the alpha prototype is not satisfactory due to low-resolution of position sensor and low control sampling rate of the PLC.

Beta version Ros-Drill as shown in Fig. 2 was designed and prototyped to be a high-precision, compact, and inexpensive setup. Essentially, servo motor is utilized to generate rotational oscillation and a closed loop control is implemented to ensure desired oscillation. Main improvements in hardware lie in microcontroller with sampling frequency of 10kHz and rotational motion sensor with resolution of 0.09° [49-51]. Despite the improvement in hardware, resolution of encoder is still very low compared with target amplitude tracking (0.2° amplitude). This is a typical mechatronic design with resource constraint, which stems from the position sensor in terms of size, resolution and costs demanded by the application [52-54]. Harmonic magnitude control is at the core of precision control and presents no particular difficulty when it uses appropriate sensors [55]. In order to track a 0.2° amplitude oscillation with sensor with 0.09° resolution, conventional control techniques fail to achieve desired tracking capability. Initially, a look-up table based gain scheduling technique was proposed to transform this low-resolution application to conventional proportional-integral-derivative (PID) method [49-50]. To counteract model uncertainty and disturbance in the control process, an adaptive control technique was designed to guarantee desired performance by adjusting control gains [56-57]. Moreover, an effective and systematic methodology which can achieve zero magnitude error tracking control for extremely low-resolution encoder was designed and implemented [58-61]. A stochastic stroke control problem was converted into a deterministic one by inferring actual motion from sensor signals [59]. The proposed control strategy consists of a two-stage adaptive control logic which adapts the control gains to achieve the actual peak-to-peak stroke. Comparisons of alpha and beta version of Ros-Drill are shown in Table 1. Controller and related peripherals are integrated into a control box.

**Fig. 2:**
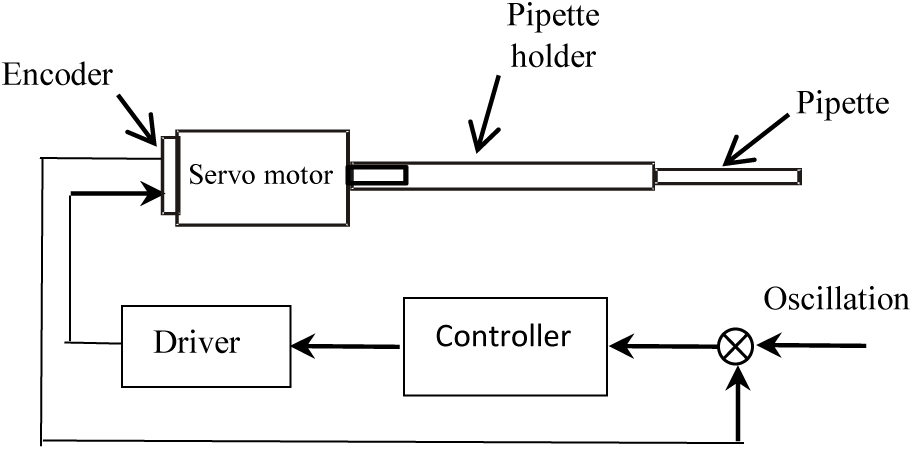
Control system of Ros-Drill setup.

**Table 1:** Comparisons of alpha and beta version of Ros-Drill.

|  | Controller | Sampling rate (kHz) | Encoder resolution (deg) | Control algorithm | Trajectory tracking capability (amplitude in deg) |
| --- | --- | --- | --- | --- | --- |
| Alpha Ros-Drill | PLC | 1 | 0.18 | PID | 1 |
| Beta Ros-Drill | Microcontroller | 10 | 0.09 | Adaptive control | 0.2 |

Ros-Drill technique looks promising from drilling mechanism and implementation perspective. Somehow, very few institutions or clinics follow this protocol and very limited experimental data is available.

### 2.4 Enhanced piezo-assisted ICSI

#### 2.4.1 Micropipettes improvement

Material and geometry of micropipettes also plays an important role in piezo-assisted ICSI. Researches and studies are conducted for optimization of geometry of micropipettes. It is shown that the Piezo-ICSI using micropipettes with a wall thickness of 0.625 μm significantly improves the survival, fertilization, good-quality day-3 embryo, pregnancy, and live birth rates when compared to the conventional ICSI and Piezo-ICSI using micropipettes with a wall thickness of 0.925 μm [62].

#### 2.4.2 Piezoelectric driven non-toxic injector

In piezo-drill cell injection, mercury is placed at the tip of pipette to reduce generation of lateral tip oscillations of injection pipette. Some alternative is proposed to suppress this excessive lateral motion in piezo-driven cell injection. A new design for piezo-actuated injector eliminates unwanted oscillations by applying physical constraints on the pipette [24]. It doesn’t rely on mercury to attenuate destructive lateral tip oscillations of the injector pipette. This mercury-free injector can sublime the piezoelectric driven injection technique to completely non-toxic level [24].

Essentially, the new injector employs a rigid attachment to limit lateral motion from piezo actuator. The efficacy of this limiter is demonstrated by the suppression of lateral movement under high speed camera by 50% [24]. However, no experimental or clinical data is available for this newly proposed method. From engineering perspective, additional constraint on actuator in one dimension usually compromises actuation capabilities of other dimensions due to mechanical coupling. Piercing force from axial oscillation of piezo-electric device is inevitably attenuated as a consequence. More simulation/modeling, e.g. finite element analysis and experiments need to be performed to demonstrate efficacy of this technique.

## 3. Methodology comparison

This section provides comparison of three major injection methods, i.e. conventional, piezo-assisted and Ros-Drill, as shown in Table 2. Conventional injection is manual and labor-intensive compared with piezo-assisted and Ros-Drill. From actuation perspective, piezo provides axial vibration whereas Ros-Drill rotational oscillation. In spite of availability of commercialized medical device for conventional ICSI, satisfactory success rate still calls for intensive training in biologists and practitioners. Advanced control algorithm and mechatronic design enable Ros-Drill to achieve desired motion at low cost, which puts Ros-Drill at a favorable position in contrast to Piezo-assisted device. The statistics for publications pertinent to injection methods of ICSI are based on data from 1995 to 2017.

**Table 2:** Comparisons between three major injection techniques for ICSI.

| | Motion type | Survival rate (%) | Fertilization rate (%) | Cost (\$k) | Drawback | # Publication (1995-2017) | # Usages |
| --- | --- | --- | --- | --- | --- | --- | --- |
| Conventional | Manual | 80 | 45 to 70 | N/A | labor intensive | >300 | 500 |
| Piezo | Axial | 90 | 75 | >5 | mercury | 85 | 300 |
| Ros-Drill | Rotational oscillation | 70 | 80 | 0.6 | insufficient data | 18 | N/A |

Nowadays, based on number of related publications and deployment, piezo-assisted method dominates in experimental and clinic activities in ICSI. In terms of survival and fertilization rate, Ros-Drill and Piezo-assisted method are comparable. However, since much more experimental data is available for Piezo-assisted method than Ros-Drill, the survival and fertilization rate for Piezo-assisted method is more statistically significant. Again Ros-Drill technique look promising from drilling mechanism and proposed implementation perspective. Somehow, very few institution or clinics follow this protocol and very limited experimental data is available.

## 4. Mechanism of membrane recovery after drilling

In essence, understanding of mechanism of membrane healing is crucial in ICSI from two-folds: a. new drilling method to facilitate membrane healing, b. method to facilitate healing after drilling.

The cell membrane, also called plasma membrane, is very thin with thickness of a few molecules. The major role of cell membrane is to isolate the intracellular contents. In addition, the cell membrane also serves to maintain the shape of the cells. Another function of the membrane is to control intercellular transport via endocytosis and exocytosis. The cargo lipids and proteins are moved from the cell membrane as substances are internalized through endocytosis whereas vesicles containing lipids and proteins fuse with the cell membrane to cargo in important proteins and enzymes. The cell membrane primarily consists of a mix of membrane proteins and lipids. The occupation of lipids can range from 20 to 80 percent of the membrane, relying on the cell type and their roles. Lipids help to maintain membranes flexibility, membrane proteins regulate the cell’s chemical environment and promote the transfer of molecules via the membrane. Phospholipids are a major component of cell membranes. Phospholipids form a bilayer where their hydrophilic head areas are oriented to the extracellular fluid, whereas their hydrophobic tail stay away from the cytosol and extracellular fluid. The lipid bilayer is semi-permeable. It allows limited molecules to diffuse through the membrane. Cholesterol is another major lipid component of cell membranes. Cholesterol molecules are largely distributed between membrane phospholipids to prevent cell membranes from stiffening through keeping phospholipids from being packed together. Glycolipids present on cell membrane surfaces with function of helping the cell to identify other cells of the body. The cell membrane contains two distinct types of associated proteins as well. Peripheral membrane proteins are exterior to the membrane. They are connected to membrane by binding with other proteins. Integral membrane proteins are inserted into the membrane, exposing on both sides of the membrane. Cell membrane proteins have various functions. Structural proteins play an essential role in the cell support and shape. Cell membrane receptor proteins is vital for cells communication with their external environment through the use of hormones, neurotransmitters, and other signaling molecules. Transport proteins of globular proteins, transport molecules across cell membranes through diffusion. Glycoproteins are embedded in the cell membrane with a carbohydrate chain attached to them, which contributes the cell communications and molecule transport across the membrane.

The membrane does not contribute to the cell’s mechanical strength, which is attributed to the cytoskeleton or cell wall. In addition to the lipid bilayer and integral proteins, RBC membrane possesses a 2D cytoskeleton tethered to the lipid bilayer. The RBC membrane cytoskeleton has a 2D six-fold structure, and is made of the spectrin tetramers connected at the actin junctional complexes, forming a 2D six-fold structure. The cytoskeleton is connected to the lipid bilayer via the “immobile” band-3 proteins at the spectrin-ankyrin binding sites and the glycophorin protein at the actin junctional complexes.

Membrane can heal after drilling. Once the micropipette is withdrawn, the lipid molecules rearrange themselves to reduce exposure of their hydrophobic regions to water. However, the rearrangement of the lipids at the pore boundary results in configurations that are not as energetically favorable as the bilayer itself, such that there is an energy penalty for the formation of holes. This energy penalty can be described by a line tension or edge tension, which is an energy per unit length along the boundary of the hole. The effective edge tension is temperature-dependent and vanishes at sufficiently high temperature, where the membrane is unstable against hole formation even in the absence of mechanical stress. Therefore, it drives the lipid molecules to the hole to reduce the line tension until the membrane is healed. We perform coarse-grained molecular dynamic simulation to illustrate the healing process, as shown in Fig.3. In the applied cell model, CG particles are introduced to represent lipid bilayer and transmembrane proteins (see Fig. 3). The CG particles carry both translational and rotational degrees of freedom (**x**_i_, **n**_i_), where **x**_i_ and **n**_i_ are the position and the orientation (direction vector) of particle *i*, respectively. The rotational degrees of freedom obey the normality condition |**n**_i_| = 1. Thus, each CG particle effectively carries 5 degrees of freedom. **x**_ij_ = **x**_j_ − **x**_i_ is defined as the distance vector between particles *i* and *j*. Correspondingly, *r*_ij_ ≡ |**x**_ij_| is the distance, and 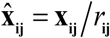 is a unit vector. The CG particles, forming the lipid membrane and membrane proteins, interact with one another via a pair-wise additive potential

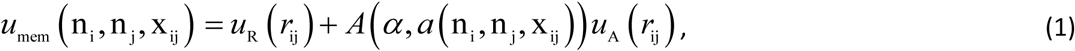

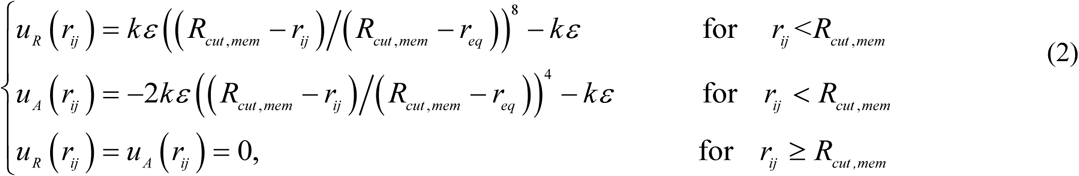

**Fig.3.**
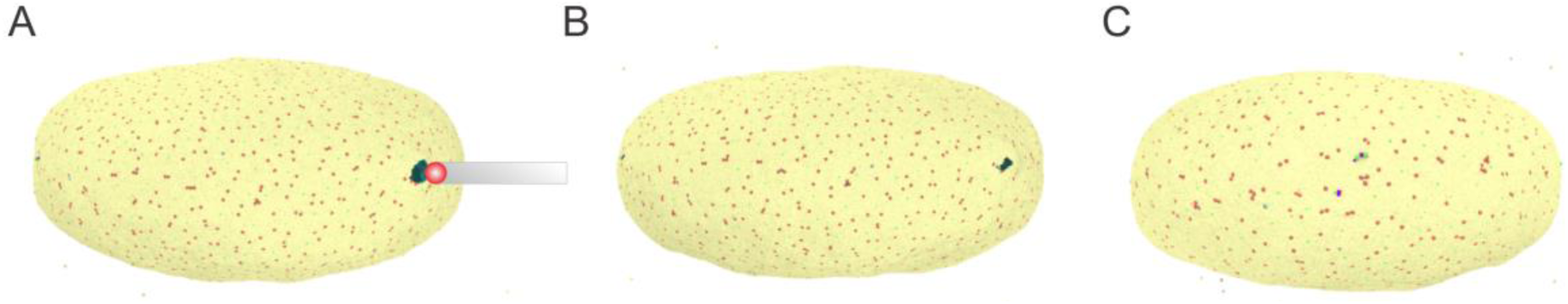
Healing of cell membrane after withdrawing the micropipette.

The healing process is highly dependent on the mechanical state of the membrane. When the membrane is under swelling (hydrated), it becomes thinner and its hydrophobic core is increasingly exposed to water so it ruptures, releasing tension through increasing the hole such that the cell becomes ruptured. On the other hand, when the membrane is under compression (dehydrated cell), the lipid molecules tend to migrate to the hole to release the compression and thus accelerate the healing the membrane. Therefore, we hypothesis that the dehydrated examined cells will facilitate healing after drilling. Our above discussions are based on the analysis of the mechanics of the lipid bilayer and cytoskeleton. We plan to validate our hypothesis by performing numerical simulation through using particle-based cell membrane and fiber models [68-77] with appropriate description of hydrodynamic interactions [78-79]. We also hope that these discussions can potentially stimulate and steer new experiments in this area.

## 5. Conclusion

A significant amount of research effort has been devoted to injection method for ICSI to achieve minimal biological damage to facilitate a rapid healing of membrane. Major milestones for injection techniques in ICSI is summarized. Their comparisons are also presented from engineering perspective. Piezo-assisted method is dominant in ICSI activities, around which further improvement in ICSI is built. Ros-Drill technique looks promising from drilling mechanism and proposed implementation perspective. However, very few institution or clinics follow this protocol and very limited experimental data is available. At last, we present a mechanism for healing process of membrane after drilling, which could potentially provide guidance for improvement of injection method.

